# Structural variation of the malaria-associated human glycophorin A-B-E region

**DOI:** 10.1101/722371

**Authors:** Sandra Louzada, Walid Algady, Eleanor Weyell, Luciana W. Zuccherato, Paulina Brajer, Faisal Almalki, Marilia O Scliar, Michel S Naslavsky, Guilherme L Yamamoto, Yeda A O Duarte, Maria Rita Passos-Bueno, Mayana Zatz, Fengtang Yang, Edward J Hollox

**Affiliations:** Wellcome Sanger Institute, Hinxton, Cambridge, UK; Department of Genetics and Genome Biology, University of Leicester, UK; Department of Parasitology, Universidade Federal de Minas Gerais, Belo Horizonte, Brazil; Human Genome and Stem Cell Research Center, Department of Genetics and Evolutionary Biology, Instituto de Biociências, Universidade de São Paulo, São Paulo, Brazil; School of Nursing, Universidade de São Paulo, São Paulo, Brazil; Laboratory of Cytogenomics and Animal Genomics (CAG), Department of Genetics and Biotechnology, University of Trás-os-Montes and Alto Douro (UTAD), Vila Real, Portugal; BioISI – Biosystems & Integrative Sciences Institute, Faculty of Sciences, University of Lisboa, Lisbon, Portugal

**Keywords:** Structural variation, copy number variation, inversion, immune response, glycophorin, *GYPA*, *GYPB*, *GYPE*, erythrocytes, malaria

## Abstract

Approximately 5% of the human genome consists of structural variants, which are enriched for genes involved in the immune response and cell-cell interactions. A well-established region of extensive structural variation is the glycophorin gene cluster, comprising three tandemly-repeated regions about 120kb in length, carrying the highly homologous genes *GYPA*, *GYPB* and *GYPE*. Glycophorin A and glycophorin B are glycoproteins present at high levels on the surface of erythrocytes, and they have been suggested to act as decoy receptors for viral pathogens. They act as receptors for invasion of a causative agent of malaria, *Plasmodium falciparum*. A particular complex structural variant (DUP4) that creates a *GYPB*/*GYPA* fusion gene is known to confer resistance to malaria. Many other structural variants exist, and remain poorly characterised. Here, we analyse sequences from 6466 genomes from across the world for structural variation at the glycophorin locus, confirming 15 variants in the 1000 Genomes project cohort, discovering 9 new variants, and characterising a selection using fibre-FISH and breakpoint mapping. We identify variants predicted to create novel fusion genes and a common inversion duplication variant at appreciable frequencies in West Africans. We show that almost all variants can be explained by unequal cross over events (non-allelic homologous recombination, NAHR) and. by comparing the structural variant breakpoints with recombination hotspot maps, show the importance of a particular meiotic recombination hotspot on structural variant formation in this region.

## Introduction

Human genetic variation encompasses single nucleotide variation, short insertion-deletions and structural variation. Structural variation includes copy number variation, tandem repeat variation, inversion and polymorphic retrotransposons. Structural variation is responsible for much of the differences in DNA sequence between individual human genomes (Sudmant et al. 2015; Zarrei et al. 2015; Hehir-Kwa et al. 2016), yet analysis of the phenotypic importance of structural variation has lagged behind the rapid progress made in studies of single nucleotide variation (Hollox and Hoh 2014; Usher and McCarroll 2015; Huddleston and Eichler 2016). This is mainly because of technical limitations in detecting, characterising, and genotyping structural variants both directly (Cantsilieris et al. 2014) and indirectly by imputation (Handsaker et al. 2015). However, a combination of new technical approaches using genome sequencing data to detect structural variation and larger datasets allowing more robust imputation of structural variation have begun to show that some structural variants at an appreciable frequency in populations do indeed contribute to clinically-important phenotypes (Sekar et al. 2016; Raffield et al. 2018).

One such example is the identification of a structural variant called DUP4 at the human glycophorin gene locus, which confers a reduced risk of severe malaria and protection against malarial anemia (Leffler et al. 2017; Algady et al. 2018; Ndila et al. 2018). The glycophorin gene locus consists of three ~120 kb tandem repeats sharing ~97% identity, each repeat carrying a closely-related glycophorin gene, starting from the centromeric end: glycophorin E (*GYPE*), glycophorin B (*GYPB*) and glycophorin A (*GYPA*) (Vignal et al. 1990; Onda et al. 1994). Large tandem repeats, like the glycophorin locus, are prone to genomic rearrangements, and indeed the DUP4 variant is a complex variant that generates a *GYPB-GYPA* fusion gene, with potential somatic variation in fusion gene copy number (Leffler et al. 2017; Algady et al. 2018). The mechanism of resistance to malaria of this gene is not fully understood, but although both glycophorin A and glycophorin B interact with receptors on *Plasmodium falciparum*, recent data suggest that alteration of receptor-ligand interactions is not important. Instead, it seems likely that DUP4 is associated with more complex alterations in the protein levels at the red blood cell surface resulting in increased red blood cell tension, mediating its protective effect against *P. falciparum* invasion (Kariuki et al. 2018). Given the size of effect of DUP4 in protection against malaria (odds ratio ~0.6) and the frequency of the allele (up to 13% in Tanzania), it is clinically very significant, although it appears to be geographically restricted to East Africa (Leffler et al. 2017; Algady et al. 2018).

Other structural variants in the glycophorin region have been identified in the 1000 Genomes project samples by using sequence read depth analysis of 1.6kb bins combined with a Hidden Markov Model approach to identify regions of copy number gain and loss (Leffler et al. 2017). This builds upon identification of extensive CNV is this area by array CGH (Conrad et al. 2009) and indeed by previous analysis of rare MNS (Miltenberger) blood groups, such as M^K^, caused by homozygous deletion of both *GYPA* and *GYPB* (Vignal et al. 1990). The variants were classified as DUP and DEL representing gain and loss of sequence read depth respectively. Although only DUP4 has been found to be robustly associated with clinical malaria phenotypes, it is possible that some of the other structural variants are also protective, but are either rare, recurrent, or both rare and recurrent, making imputation from flanking SNP haplotypes and genetic association with clinical phenotypes challenging.

It is important, therefore, to extend the catalogue of structural variants at this locus and robustly characterise their nature and likely effect on the number of full-length and fusion glycophorin genes. In this study we use sequence read depth analysis of 6466 genomes from across the world, followed by direct analysis of structural variants using fibre-FISH and breakpoint mapping using paralogue-specific PCR and Sanger sequencing. This will allow future development of robust yet simple PCR-based assays for each structural variant and detailed analysis of the phenotypic consequences of particular structural variants on malaria infection and other traits. We also examine the pattern of structural variation breakpoints in relation to their mechanism of generation and known meiotic recombination hotspots within the region, and the relative allele frequencies across the world. Together, this allows us to gain some insight into the evolutionary context of the extensive structural variation at the glycophorin locus.

## Methods

### Sequencing data

Sequence alignment files (.bam format7) from four cohorts (1000 Genomes Project ENA accession number PRJNA262923) with a mean coverage of 7.4x (Auton et al. 2015), Simons Diversity Project ENA accession number PRJEB9586 with a mean coverage of 43x (Mallick et al. 2016), and the Gambian Genome Diversity project mean coverage 4x, ENA study IDs ERP001420, ERP001781, ERP002150, ERP002385) (Band et al. 2019) were downloaded from the European Nucleotide Archive or from the International Genome Sample Resource site http://www.internationalgenome.org/data-portal/ (Clarke et al. 2017). Brazilian sequence alignment files from the SABE (Health, Wellbeing and Aging) study (Barbosa et al. 2005) and a sample of cognitively healthy octogenarians enrolled at the Human Genome and Stem Cell Research Center (80+), with a mean coverage of 30x for 1324 individuals generated at Human Longevity Inc. (HLI, San Diego, California) (Telenti et al. 2016).

DNA sequences from the 1000 Genomes project and the Simons diversity project had been previously aligned to reference GRCh37 (hg19) to generate the alignment bam files. The exception is sample NA18605, which was previously sequenced at high coverage (Lan et al. 2017) downloaded as paired-end Illumina sequences in fastq format (ENA sample accession number SAMN00001619), and aligned to GRCh37 using standard approaches: FastQC v0.11.5 and Cutadapt v01.11 to trim reads and adapters, mapping using BWA-MEM v0.7.15, processing of the BAM files using SAMtools v1.8, local realignment was done using GATK v3.6 and duplicate reads marked using Picard v.1 and removed using SAMtools. Samples from the Brazilian genomes and the Gambian genome diversity project had been aligned to GRCh38.

Throughout this paper, all loci are given using GRCh37 reference genome coordinates. For analyses on GRCh38 alignments, genome coordinates were translated from the GRCh37 coordinates using the Liftover tool within the UCSC Genome Browser (Kent et al. 2002).

### Structural variant detection

For each sample, we used SAMtools (SAMtools view ‒c ‒F 4) (Li et al. 2009) on indexed bam files to count mapped reads to the glycophorin region (chr4:144745739-145069133) and a reference region chr4:145516270-145842585. The reference region has no segmental duplications, and is absent from copy number variation according to the gold standard track of the database of Genomic Variants (DGV) (MacDonald et al. 2014). A ratio of the number or reads mapping to the glycophorin region to the number of reads mapping to the reference region allows an estimate of the total increase or decrease of sequence depth spanning the glycophorin region (reflecting copy number gain or copy number loss, respectively). Following plotting these data for each cohort on a histogram and observation of distinct clusters (supplementary figure 1), samples with a ratio below 0.9 were classified as potential deletions and those above 1.1 potential duplications.

For the samples with potential deletions and duplications, number of mapped reads was calculated across the glycophorin region in 5kb non-overlapping windows, and values, normalised to average read count and diploid copy number, were plotted. Presence and nature of structural variants were assessed by examination of the plots, and particular variants called by plotting together with a reference sample for that variant. For the Simons Diversity Project samples, 114 potential deletions were identified, much more than in other cohorts (Supplementary figure 1). Inspection of these plots showed that 101 of these samples showed a small apparent ~15kb deletion at the *GYPE* gene. This deletion was not found previously by others (Leffler et al. 2017) or by us in any other cohort, and coincides with a region of low mappability, suggesting that this may be an artefact caused either by particular filtering conditions or the particular genome assembly (GRCh37d5) that includes decoy sequences. These 101 samples were treated as being homozygous for the reference structure.

### Fiber-FISH

The probes used in this study included four WIBR-2 fosmid clones selected from the UCSC Genome Browser GRCh37/hg19 assembly and a 3632-bp PCR product that is specific for the glycophorin E repeat (Algady et al. 2018). Probes were made by amplification with GenomePlex Whole Genome Amplification Kits (Sigma-Aldrich) as described previously (Gribble et al. 2013). Briefly, the purified fosmid DNA and the PCR product were amplified and then labeled as follow: G248P86579F1, G248P89366H1 and glycophorin E repeat-specific PCR product were labeled with digoxigenin-11-dUTP, G248P8211G10 was labeled with biotin-16-dUTP, G248P85804F12 was labeled with DNP-11-dUTP and G248P80757F7 was labeled with Cy5-dUTP. All labeled dUTPs were purchased from Jena Bioscience.

The preparation of single-molecule DNA fibers by molecular combing and fiber-FISH was as previously published (Louzada et al. 2017), with the exception of post-hybridization washes, which consisted of three 5-min washes in 2× SSC at 42°C, instead of two 20-min washes in 50% formamide/50% 2× SSC at room temperature.

### Breakpoint analysis using PCR and Sanger sequencing

Using the 5kb window sequence read count data, PCR primers were designed so that a PCR product spanned the predicted breakpoints for each deletion and duplication. The 3’ nucleotide for each PCR primer was designed to match uniquely to a particular glycophorin repeat, and to mismatch the other two glycophorin repeats. Annealing specificity of the PCR primer was enhanced by incorporating a locked nucleic acid at that particular 3’ position of the PCR primer (Latorra et al. 2003). Long-range PCR amplification used 10 ng genomic DNA in a final volume of 25.5 μl, including 0.5 μl of each 10μM primer, 0.075U *Pfu* DNA polymerase, 0.625U *Taq* DNA polymerase, and 2.25 μl of PCR buffer (45 mM Tris-HCl (pH 8.8), 11 mM ammonium sulphate, 4.5 mM magnesium chloride, 6.7 mM 2-mercaptoethanol, 4.4 mM EDTA (pH 8.0), 113 μg/mL non-acetylated Bovine Serum Albumin (BSA) (Ambion®) and 1 mM of each dNTP (Promega) (Jeffreys et al. 1990)). The reaction was thermal cycled as follows: 94°C 1 minute, followed by 20 cycles of 94°C for 15s, x°C for 10 minutes, followed by 15 cycles of 94°C for 15s, x°C for 10 minutes+15s for each successive cycle, followed by a final extension at 72°C for 10 minutes, where x is the annealing temperature for a particular primer pair shown in supplementary table 1. PCR products were purified using agarose gel electrophoresis (Ma and Difazio 2008) and Sanger sequenced using standard approaches. PCR primers are shown in supplementary table 1. Multiple alignments with paralogous reference sequences used MAFFT v7 (Katoh and Standley 2013) available at the EMBL-EBI Job Dispatcher framework (Li et al. 2015). A breakpoint was called in the transition region between three paralogous sequence variants corresponding to one glycophorin repeat and three paralogous sequence variants corresponding to the alternative glycophorin repeat.

### Breakpoint analysis using high depth sequences

For particular variants, copy number breakpoints were refined by inspecting sequence read depth in 1kb windows spanning the likely breakpoints identified by the 5kb window analysis. Changes in read depth were then confirmed directly using the Integrative Genome Viewer (Thorvaldsdóttir et al. 2012).

### Nomenclature of variants

We used the same nomenclature as Leffler et al. 2017 when our variant could be identified as the same variant in the same sample from the 1000 Genomes project. In some instances, we could not distinguish particular singleton variants called from more common called variants. For example, DUP27 carried by sample NA12249 could not easily be distinguished from the more frequent DUP2, and DUP24 carried by HG04038 could not be distinguished from DUP8. Other variants, which either had not been unambiguously identified in the 1000 Genomes previously or were identified in other sample cohorts, were given DEL or DUP numbers following on from variants catalogued previously.

### Analysis of recombination hotspots

Previously published data on hotspot location and type (Pratto et al. 2014) was converted to BED format and intersected with the breakpoint locations in BED format using BEDTools v 2.28.0 (Quinlan and Hall 2010). The statistical significance of the overlap was calculated using the fisher command in BEDTools, which uses a Fisher’s exact test on the number of overlaps observed between two BED files.

## Results

### Structural variation using sequence read depth analysis

Previous work by us and others has shown that unbalanced structural variation - that is, variation that causes a copy number change - can be effectively discovered by measuring the relative depth of sequence reads across the glycophorin region (Leffler et al. 2017; Algady et al. 2018). We analysed a total of 6466 genomes from four datasets spanning the globe - the 1000 Genomes phase 3 project set, the Gambian variation project, the Simons diversity project, and the Brazilian genomes project. We took a different sequence read depth approach to that previously used, counting the reads that map to the glycophorin repeat region and dividing by the number of reads mapping to a nearby non-structurally variable region to normalise for read depth. By analysing each cohort of diploid genomes as a group, we could identify outliers where a higher value indicated a potential duplication or more complex gain of sequence, and lower values indicated a potential deletion (Supplementary figure 1). Sequence read depth was analysed in 5kb windows across each of the outlying diploid genomes to identify and classify the structural variant.

Since structural variant calling had been previously done on the 1000 Genomes project cohort, this provided a useful comparison to assess our approach. We analysed 2490 samples from this cohort and identified five distinct deletions carried by 88 individuals that were previously identified, and 16 distinct duplications carried by 34 individuals (table 1) that were also previously identified. We also identified a new duplication variants, termed DUP29 (a duplication of GYPB), that had not been robustly identified previously in that cohort. However, as expected, smaller duplications, most notably DUP1, were not detected by our approach. We extended our analysis to 390 Gambian diploid genomes and identified 51 samples with DEL1 or DEL2 variants, and DEL16, subsequently characterized in the Brazilian cohort below. Two samples were heterozygous for the duplication DUP5.

**Table 1.**
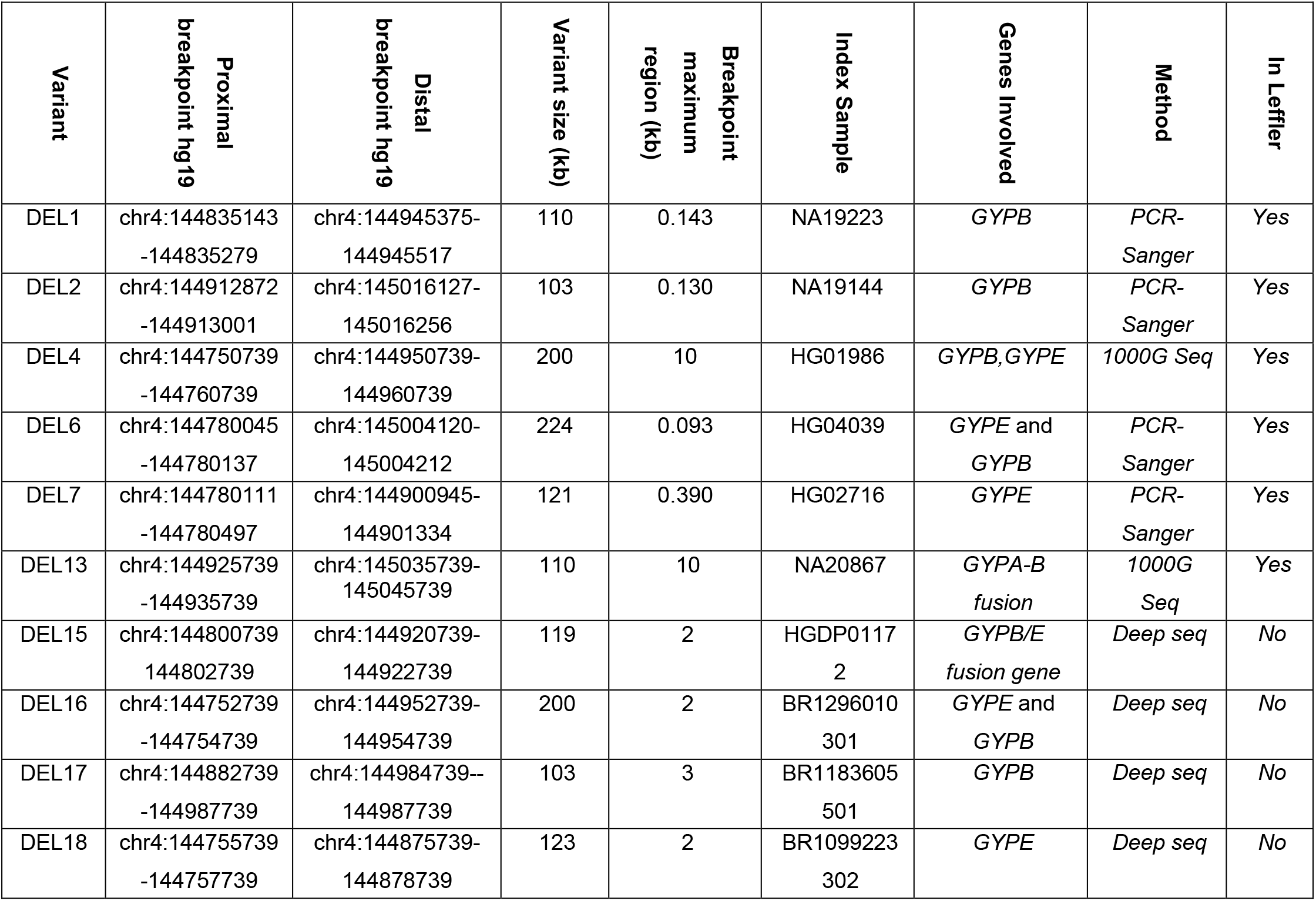
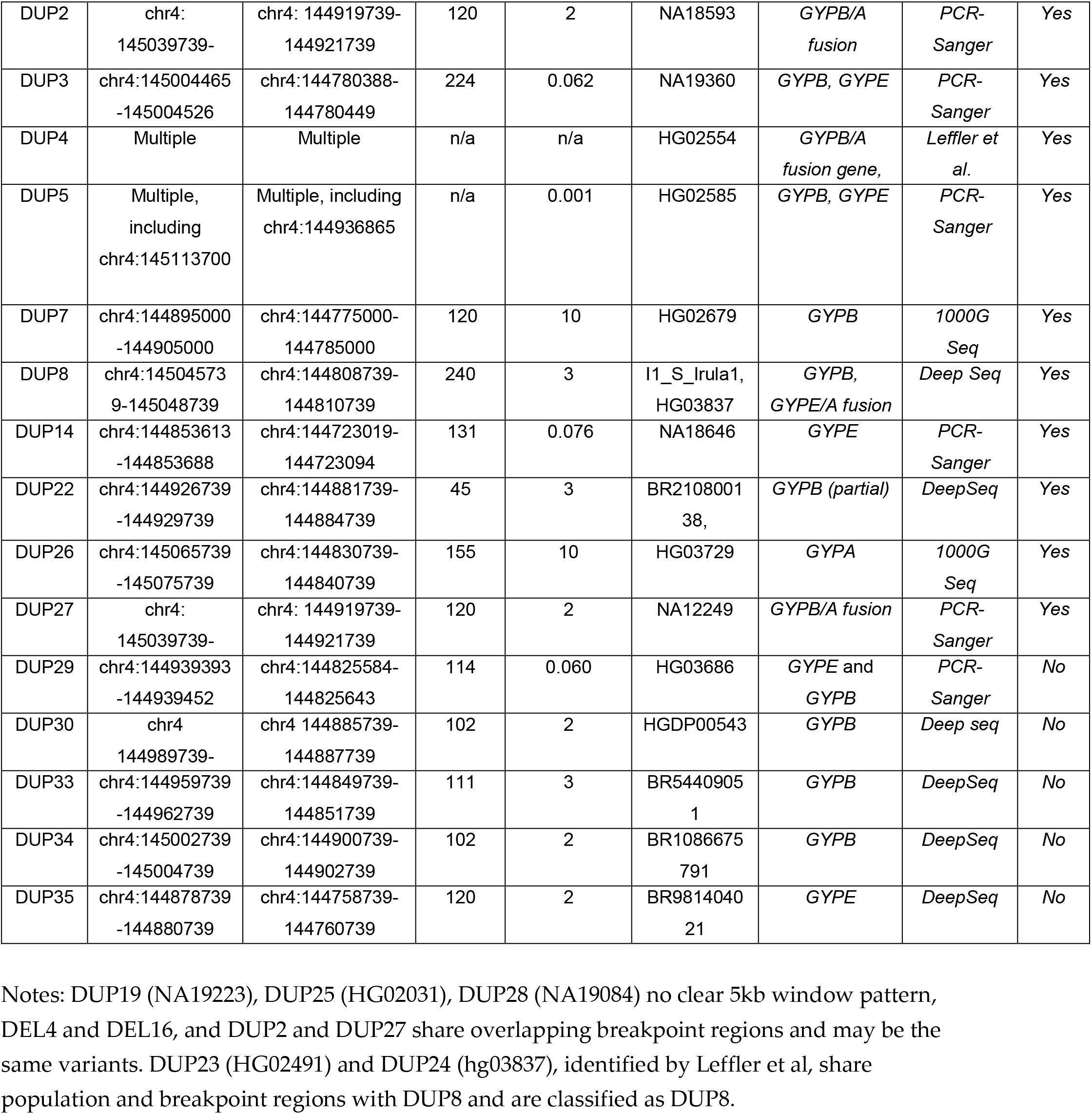
Glycophorin structural variants identified in this study.

**Table 2.** Frequency of structural variants observed more than once across the cohorts analysed.

|  | 1000 Genomes |  |  |  |  | Gambian | Simons | Brazilian |
| --- | --- | --- | --- | --- | --- | --- | --- | --- |
| Population | EUR | AFR | SAS | EAS | AMR | AFR | ALL | AMR |
| Total chromosomes | 600 | 640 | 386 | 606 | 258 | 782 | 546 | 2648 |
| DEL1 | 0 | 53 | 0 | 1 | 1 | 55 | 7 | 19 |
| DEL2 | 0 | 26 | 0 | 0 | 2 | 2 | 4 | 12 |
| DEL4/16 | 0 | 1 | 0 | 0 | 0 | 1 | 0 | 3 |
| DEL6 | 0 | 0 | 1 | 0 | 1 | 0 | 0 | 0 |
| DEL7 | 0 | 2 | 0 | 0 | 0 | 0 | 1 | 0 |
| DUP2/27 | 0 | 1 | 1 | 11 | 1 | 0 | 0 | 7 |
| DUP3 | 0 | 4 | 0 | 0 | 0 | 0 | 0 | 0 |
| DUP5 | 0 | 1 | 0 | 0 | 0 | 2 | 0 | 0 |
| DUP7 | 0 | 1 | 0 | 0 | 1 | 0 | 0 | 2 |
| DUP8 | 0 | 0 | 4 | 0 | 0 | 0 | 1 | 2 |
| DUP29 | 0 | 0 | 1 | 0 | 0 | 0 | 1 | 0 |
| DUP22 | 0 | 0 | 0 | 1 | 0 | 0 | 1 | 1 |
| DUP30 | 0 | 0 | 0 | 0 | 0 | 0 | 3 | 0 |
| DUP35 | 0 | 0 | 0 | 0 | 0 | 0 | 0 | 2 |

Both 1000 Genomes and Gambian Genome Diversity samples have been sequenced to low depth. High depth sequencing will allow more robust identification of structural variation by improving the signal/noise ratio of sequence read depth analysis. We analysed the publically available high-depth data from the Simons Diversity Project for glycophorin variation. From the 273 individuals, 4 different deletion types were carried by 13 individuals, and 3 different duplication types were carried by 5 individuals. A novel deletion, DEL15 was identified which deleted part of *GYPB* and part of *GYPE* in an individual from Bergamo in Italy, and a novel duplication was observed in three individuals from Papua New Guinea, termed DUP30 and duplicating the *GYPB* gene. Another duplication variant (DUP8), which is the largest variant found so far, duplicated 240kb, creating an extra full length *GYPB* gene and a *GYPE-GYPA* fusion gene (Table 1).

A further 1324 samples sequenced to high coverage diploid genomes from Brazil were analysed, which, given the extensive admixture from Africa in the Brazilian population, are likely to be enriched for glycophorin variants from Africa. Four new duplication variants (DUP33-DUP35) and three new deletion variants were found (DEL16, DEL17, DEL18), two of which of which delete the *GYPB* gene (Table 1).

### Fibre-FISH analysis of structural variants

Sequence read depth analysis shows copy number gain and loss with respect to the reference genome to which the sequence reads are mapped, but it does not determine the physical structure of the structural variant. For the more common glycophorin structural variants, we used fibre-FISH in order to determine the physical structure. In all cases, a set of multiplex FISH probes, with each probe being visualised with a unique fluorochrome, was used so that the orientation and placement of the repeats could be identified (Figure 1). The repeated nature of the glycophorin region means that the green and red probes from the *GYPB* repeat cross-hybridise with the other repeats, with the *GYPA* repeat distinguishable from the *GYPB* and *GYPE* repeats by a 16kb insertion resulting in a small gap of signal in the green probe (Figure 1).

**Figure 1.**
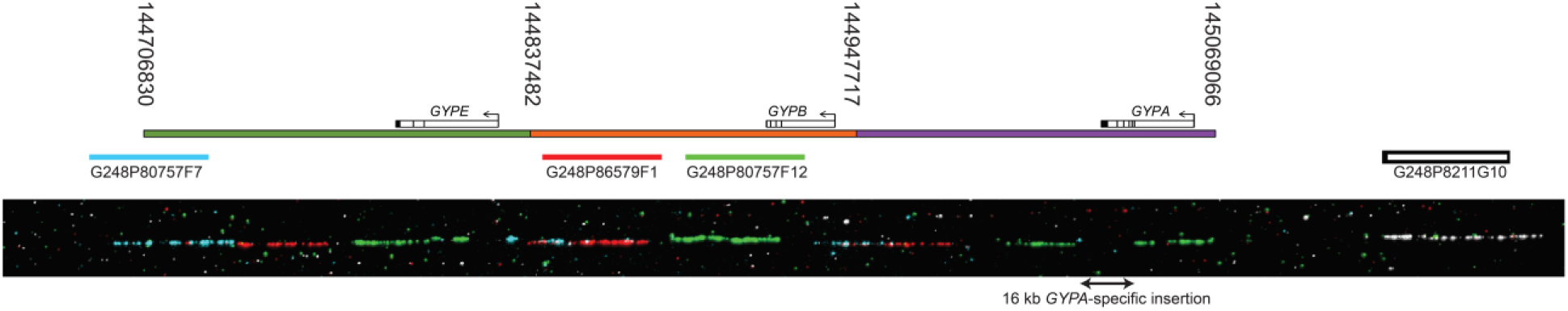
Structure of the glycophorin reference allele. A representation of the reference allele assembled in the GRCh37/hg19 assembly is shown, with the three distinct paralogous ~120kb repeats of the glycophorin region coloured green, orange and purple, carrying *GYPE*, *GYPB* and *GYPA* respectively. Numbers over the start and end of each paralogue represent coordinates in chromosome 4 GRCh37/hg19 assembly. Coloured bars represent fosmids used as probes in fibre-FISH, with the fosmid identification number underneath. An example fibre FISH image of this reference haplotype (from sample HG02585) is shown below.

For most variants the fibre-FISH results confirmed the structure previously predicted (Leffler et al. 2017) and expected if the variants had been generated by non-allelic homologous recombination between the glycophorin repeats (Figures 2 and 3). However, three variants showed a complex structure that could not be easily predicted from the sequence read depth analysis. The DUP4 variant shows a complex structure and has been described previously (Algady et al. 2018). Two other structural variants (DUP5 and DUP26) also showed complex patterns of gains or losses, and fibre-FISH clearly shows the physical structure of the variant, including inversions.

**Figure 2.**
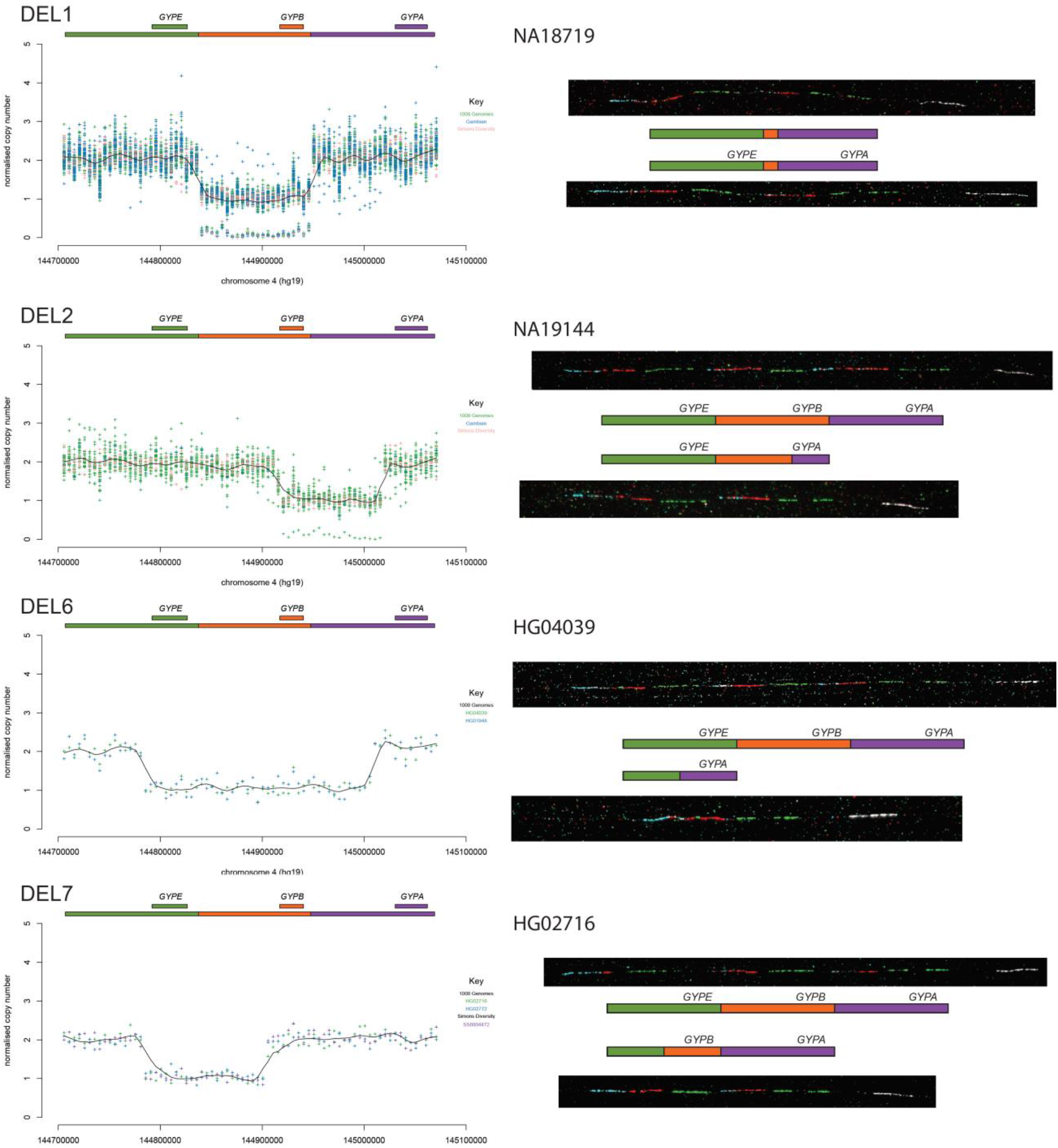
Fibre-FISH validation of four glycophorin deletions. Sequence read depth (SRD) analysis of selected deletions (DEL1, DEL2, DEL6, DEL7) is shown on the left, with sample, or cohort, coloured according to the key on each plot, together with a Loess best-fit line Note for DEL1 and DEL2 homozygous individuals are detected with a normalised copy number of zero across the deletion. Representative fibre-FISH images from the index sample of each variant are shown on the right, with clones and fluorescent labels as shown in figure 1. All index samples apart from NA18719 are heterozygous, with a representative reference (top) and variant (bottom) allele from that sample shown. A schematic diagram next to the corresponding fibre-FISH image shows the structure of each allele inferred from the fibre-FISH and SRD analysis.

**Figure 3.**
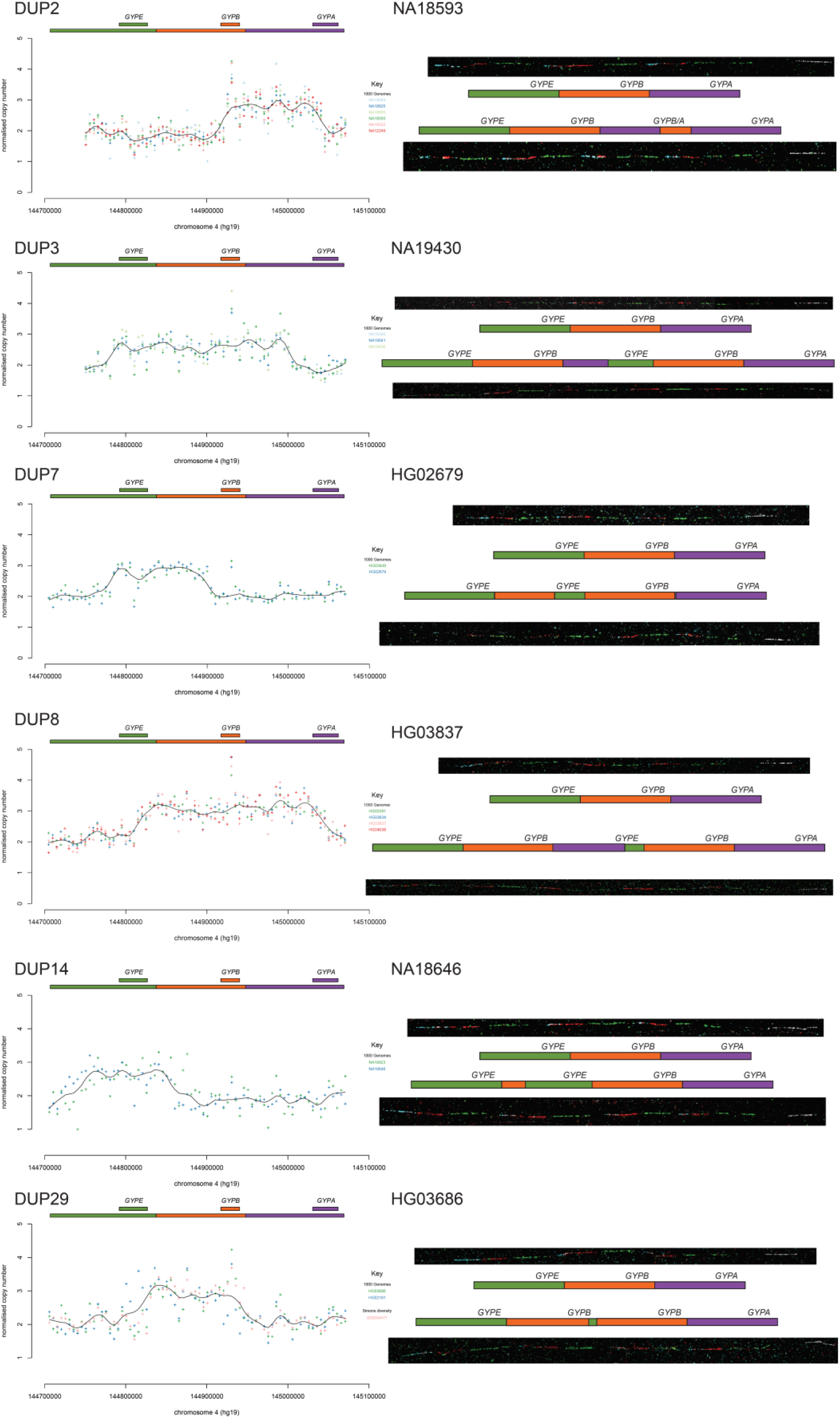
Fibre-FISH validation of six glycophorin duplications. Sequence read depth (SRD) analysis of selected duplications (DUP2, DUP3, DUP7, DUP8, DUP14 and DUP29) is shown on the left, with sample, or cohort, coloured according to the key on each plot, together with a Loess best-fit line. Representative fibre-FISH images from the index sample of each variant are shown on the right, with clones and fluorescent labels as shown in figure 1, (with an additional green-labelled PCR product specific to the glycophorin E repeat for HG03686). All index samples are heterozygous, with a representative reference and variant allele from that sample shown. A schematic diagram next to the corresponding fibre-FISH image shows the structure of each allele inferred from the fibre-FISH and SRD analysis.

The more frequent of these two complex structural variants, DUP5, seems to be restricted to Gambia, as it is found once in the GWD population from the 1000 Genomes project and twice in the Jola population from the Gambian Genome diversity project (Table 1). Sequence read depth analysis suggests that DUP5 has two extra copies of *GYPE* and an extra copy of *GYPB*, with an additional duplication distal of *GYPA* outside the glycophorin repeated region (Figure 4a). Fibre-FISH analysis on cells from an individual carrying the DUP5 variant (HG02585) confirmed heterozygosity of the variant, with one allele being the reference allele, and revealed, for the first time, that the variant allele presents a complex pattern of duplication and rearrangement, with part of the fosmid (pseudocoloured in white) mapping distal to *GYPA* being translocated into the glycophorin repeated region, adjacent to the green-coloured fosmid (Figure 4b). Alternative fibre-FISH analysis using an additional fosmid probe mapping distally, and labelled in red, confirmed this (Figure 4c). The pattern of FISH signals occurring distally to the translocation suggests that the immediately adjacent glycophorin repeat is inverted. To distinguish the distal end of the *GYPB* repeat from the distal end of the *GYPE* repeat, a pink-coloured probe from a short *GYPE*-repeat-specific PCR product was also used for fibre-FISH, and clearly shows only a single copy of the distal end of the *GYPB* repeat in the DUP5 variant, at the same position as the reference. The predicted breakpoint between the non-duplicated sequence distal to *GYPA* and duplicated sequence within the duplicated region was amplified by PCR and Sanger sequenced, confirming that the non-duplicated sequence was fused to an inverted *GYPB* repeat sequence (Figure 4d). The model suggested by the fibre-FISH and breakpoint analysis is consistent with the overall pattern of sequence depth changes observed (Figure 4a). The sequence outside the glycophorin repeat corresponds to an ERV-MaLR long terminal retroviral element, but the sequence inside the glycophorin repeat sequence is not, suggesting that non-allelic homologous recombination was not the mechanism for formation of this breakpoint. However, there is a 4bp microhomology (GTGT) between the two sequences, suggesting that microhomology-mediated end joining could be a mechanism for formation of this variant.

**Figure 4.**
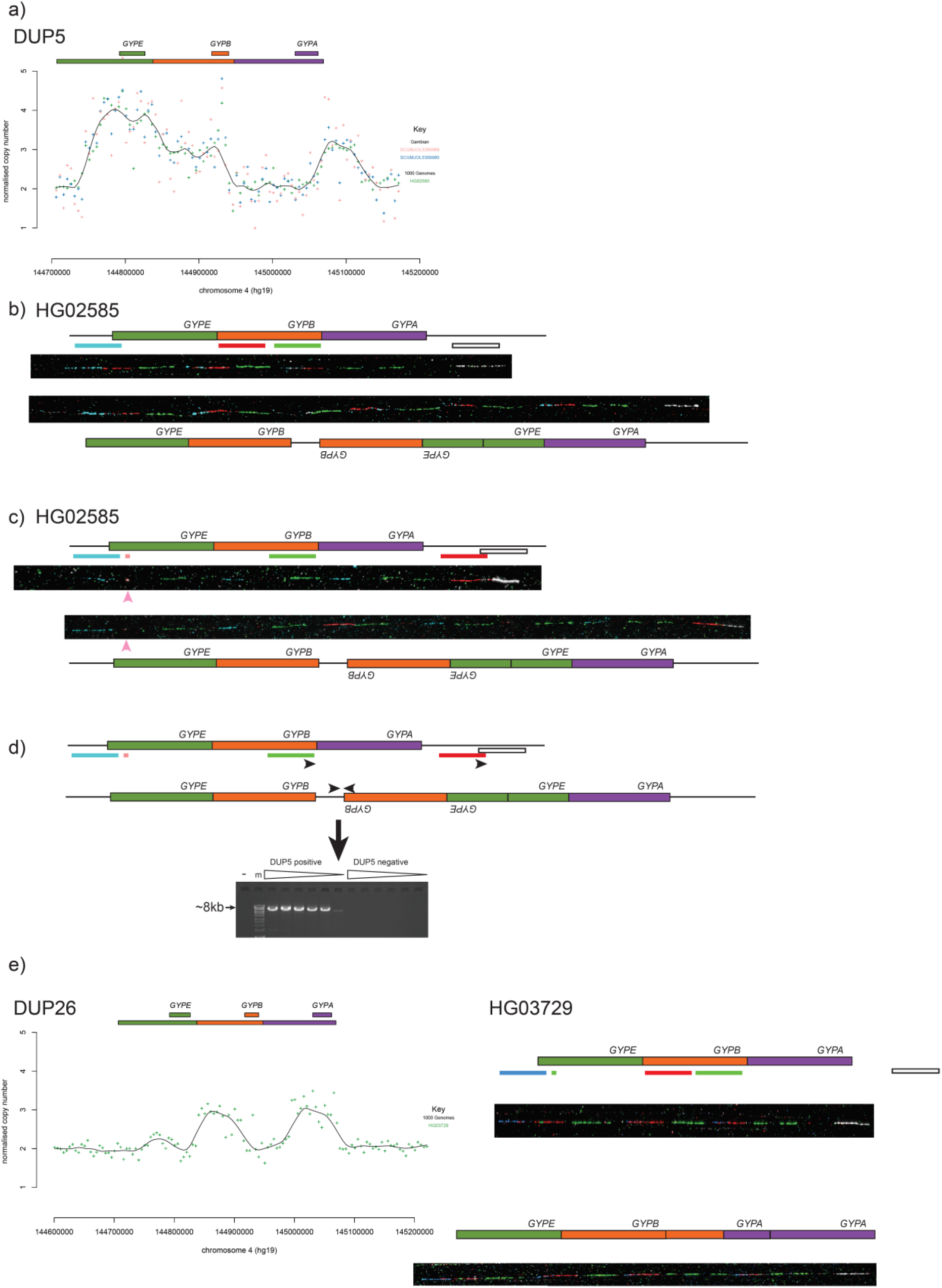
Analysis of DUP5 and DUP26 complex structures. a) Sequence read depth (SRD) analysis of three individuals heterozygous for the DUP5 variant. b) Representative fibre-FISH images from the DUP5 index sample HG02585. Clones and fluorescent labels as shown in figure 1. c) Representative fibre-FISH images from the DUP5 index sample HG02585. Clones and fluorescent labels as shown in figure 1, except the red probe is fosmid G248P89366H1 and the pink probe is the glycophorin E repeat-specific PCR product. d) Schematic showing design of PCR primers for specific amplification (black arrows) on reference and DUP5 structures. The ethidium bromide stained agarose gel shows a ~8kb PCR product generated by these DUP5 specific primers. HG02554 is the DUP5 sample, “-“ indicates a negative control with no genomic DNA and the marker, indicated by “m”, is Bioline Hyperladder 1kb+. The triangles indicate increasing PCR annealing temperature from 65°C to 67°C. e) Sequence read depth (SRD) analysis (left) and fibre-FISH analysis (right) of the index sample HG03729 heterozygous for DUP26 variant. Fosmid clones for fibre-FISH are as figure 1, except with the addition of the glycophorin E repeat-specific PCR product labelled in green.

The DUP26 variant was observed once, in sample HG03729, an Indian Telugu individual from the United Kingdom, sequenced as part of the 1000 Genomes project. Sequence read depth analysis predicts an extra copy of the glycophorin repeat, partly derived from the *GYPB* repeat and partly from the *GYPA* repeat (Figure 4e). The fibre-FISH shows an extra repeat element that is *GYPB*-like at the proximal end and *GYPA*-like at the distal end, and carries the *GYPA* gene. This structure is unlikely to have been generated by a straightforward single NAHR event, and we were unable to resolve the breakpoint at high resolution.

### Breakpoint analysis of structural variants

Defining the precise breakpoint of the variants can allow a more accurate prediction of potential phenotypic effects of each variant by assessing, for example, whether a glycophorin fusion gene is formed or whether key regulatory sequences are deleted. We used two approaches to define breakpoints. The first approach identified the two 5kb windows that spanned the change in sequence read depth at both ends of the deletion or duplication, and by designing PCR primers to specifically amplify across the junction fragment (Figure 5a,b), variant-specific PCR amplification produces an amplicon that can be sequenced (Figure 5c). After Sanger sequencing the amplicons, the breakpoint can be shown to be where a switch occurs between paralogous sequence variants (PSVs) that map to different glycophorin repeats (Figure 5d), supporting the model that a NAHR mechanism is responsible for generating these structural variants (Figure 5e). The second approach makes use of high depth sequencing. The two 5kb windows spanning the change in sequence read depth are again identified and sequence read depth calculated in 1kb windows to further refine the breakpoint. The sequence alignment spanning the two 1kb windows is examined manually for paired sequence reads where the gap between the aligned pairs is consistent with the size of the variant, or where both sequence pairs align but one aligns with multiple sequence mismatches.

**Figure 5.**
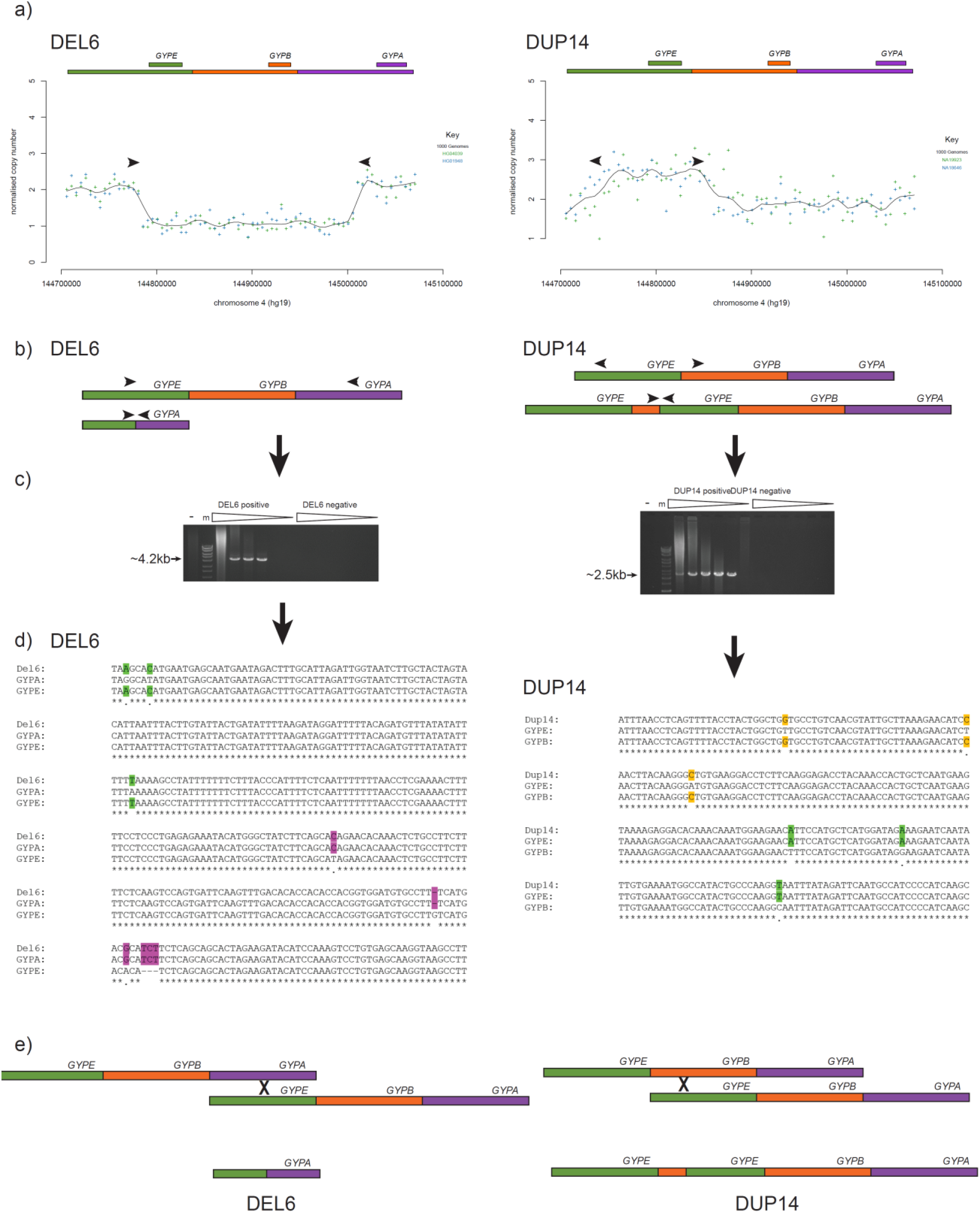
Examples of refining breakpoints of a deletion (DEL6) and a duplication (DUP14) a) Sequence read depth analysis, indicating position of PCR primers (not to scale) b) Variant model, showing position of primers on reference and variant c) Agarose electrophoresis of long PCR products using variant-specific primers indicated in b). “-“ indicates a negative control with no genomic DNA and the marker, indicated by “m”, is Bioline Hyperladder 1kb+. The triangles indicate increasing PCR annealing temperature from 58°C to 67°C. d) Multiple sequence alignment of the variant-specific PCR product, with homologous sequence on the *GYPA* repeat and the *GYPE* repeat. *GYPE*-specific variants are in green, *GYPA*-repeat-specific variants are in purple. e) A model of the generation of the variants by NAHR.

With the exception of DEL4, DUP7 and DUP26, where only low-coverage sequence was available, all other breakpoints could be localised to between 10 kb and 1 bp. For most variants, the breakpoints occur between genes resulting in loss or gain of whole genes, and therefore likely to show gene dosage effect. It is known that DUP4 results in a *GYPB-GYPA* fusion gene that codes for the Dantu blood group, and a fusion gene is also predicted for DUP2, DUP8 and DEL15. The DUP2 variant generates a *GYPB-GYPA* fusion gene comprising exons 1-2 of *GYPB* and exons 4-7 of *GYPA* corresponding to the St^a^ (GP.Sch) blood group (Anstee et al. 1982; Daniels 2008). Breakpoint analysis of NA12249, the sample carrying the DUP27 variant, showed that DUP27 breakpoint is in the same intron as DUP2, although the exact breakpoint is complex and it is unclear whether DUP27 is exactly the same variant as DUP2.

The DUP8 variant is predicted to generate a fusion gene consisting of exon 1 of *GYPE* and exons 2-7 of *GYPA*, and the DEL15 variant is predicted to combine the first two exons of *GYPB* with the final three exons of *GYPE*. It is unlikely that DUP8 has a phenotype, given the involvement of the 5’ end of *GYPE*, which is not expressed. However, the DEL15 variant is predicted to generate a *GYPB-GYPE* peptide, similar to the rare U- blood group phenotype which has a breakpoint between exon 1 of *GYPB* and exon 2 of *GYPE*, resulting in a lack of expression of glycophorin B in homozygotes (Rahuel et al. 1991). Other variants involve breakpoints within 1kb of a gene coding region and could potentially affect expression levels of the neighbouring gene.

### Mechanism of formation of structural variants

The pattern of deletions and duplications observed is consistent with a simple NAHR mechanism of formation for the variants (Figure 5e), with the exception of DUP5 and DUP26. We investigated whether the breakpoints we had found co-localised with known meiotic recombination hotspots previously determined by anti-DMC1 ChIP-Seq of the testes of five males (Pratto et al. 2014). Importantly, the recombination hotspot dataset mapped hotspots in individuals carrying different alleles of the highly-variable PRDM9 protein, a key determinant of recombination hotspot activity, with different alleles activating different recombination hotspots. The glycophorin region contains one hotspot regulated by the PRDM9 A allele, common in Europeans (allele frequency 0.84), and the PRDM9 C allele, common in sub-Saharan Africans (allele frequency 0.13). In our data we found no breakpoints coincident with the PRDM9 A allele hotspot but 4 breakpoints coincident with the PRDM9 C allele hotspot (Figure 6). The overlap between the PRDM9 C allele hotspot and the structural variant breakpoints is statistically significant (two-tailed Fisher’s exact test, p=0.012) and reflects the observation that there are more different rare structural variants in sub-Saharan African populations, with high frequencies of the C allele, than in European populations where the C allele is almost absent (allele frequency 0.01) (Berg et al. 2011).

**Figure 6.**
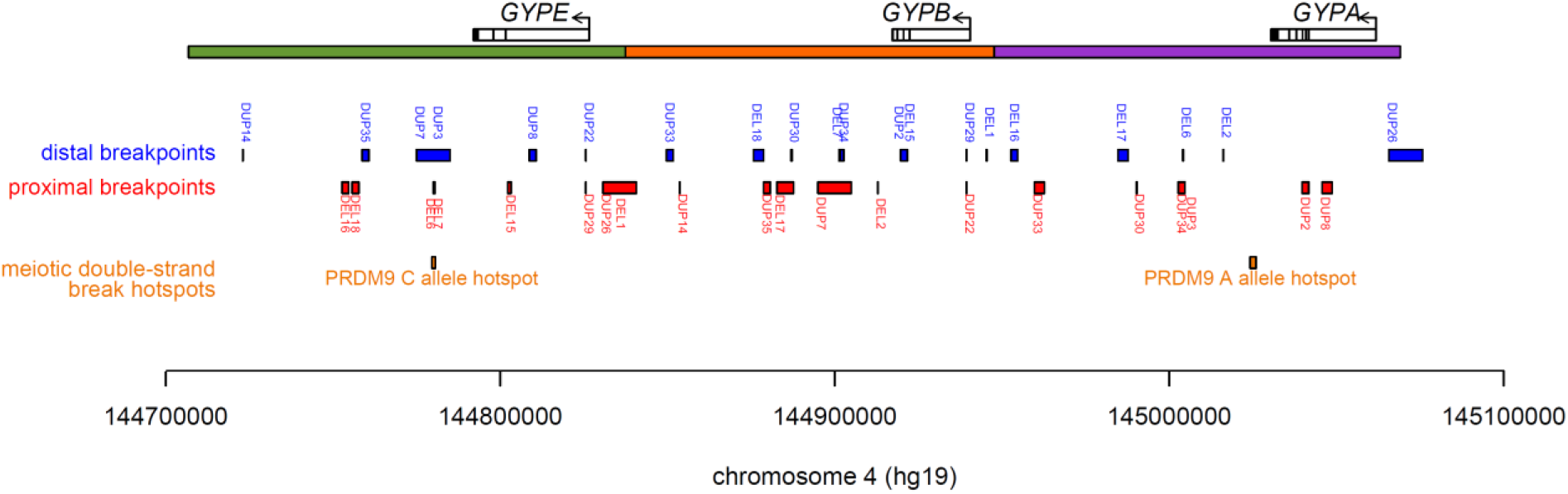
Structural variant breakpoints and meiotic recombination hotspots. The glycophorin region is shown together with the glycophorin genes. Below are the breakpoint regions for each structural variant, labelled in blue for the distal breakpoint in the variant, and red for the proximal breakpoint in the variant. Meiotic double strand break hotspots, corresponding to recombination hotspots (Pratto et al. 2014) are shown in orange, labelled the PRDM9 allele responsible for activating that hotspot.

## Discussion

We have characterised a number of structural variants at the human glycophorin locus. These are almost always large deletions or duplications involving the loss or gain of one or occasionally two glycophorin repeat regions of about 120kb. These losses and gains have been generated by non-allelic homologous recombination (NAHR) mutational events, with particular involvement of the PRDM9 C allele, which is at appreciable frequencies in African populations and directs high recombination rates at its cognate recombination hotspots. A more complex variant, termed DUP5, was also characterised, and was shown to be an inversion-duplication generated by at least 1 microhomology-mediated end-joining event involving DNA sequence outside the glycophorin repeat region. Similarly, DUP26 is a complex variant that is unlikely to have been generated by a single NAHR event.

Only DEL1, DEL2 and DUP2 are frequent variants. Both DEL1 and DEL2 delete the *GYPB* gene and it is tempting to speculate that their high frequency in African populations and populations with African admixture is due to selection. However, the absence of evidence for any protective effect against malaria argues against malaria being the cause of this selection, so this remains speculation. DUP2 is at notable frequencies particularly in East Asia, and is predicted to generate a *GYPB-GYPA* fusion gene corresponding to the St^a^ blood group, which is known to be at appreciable frequencies in East Asia (Madden et al. 1964). In this region, malaria infections are caused by *Plasmodium falciparum* as well as *Plasmodium vivax*; alternatively, this fusion gene may facilitate glycophorins acting as a decoy receptor for other pathogens, such as hepatitis A virus (Sanchez et al. 2004). Previous work suggests that DUP2 has arisen on multiple haplotype backgrounds (Leffler et al. 2017), which suggests a large East Asian population panel is need for future accurate imputation.

Other variants seem either to be geographically localised (for example DUP5) or very rare and detected as singletons in our dataset. This suggests that analysis of other large genomic datasets will discover further unique glycophorin structural variants, and that much glycophorin structural variation is individually rare but collectively more frequent, leading to challenges in imputing glycophorin SV from SNP GWAS data.

In contrast to other studies, we used a three-step approach to determine copy number. We used read counts over the whole glycophorin region to detect samples with duplications (more than expected number of reads) and deletions (fewer than expected number of reads). We then used window-based analysis of sequence read depth and paralogue-specific allele-specific PCR and Sanger sequencing to refine copy number breakpoints. Finally, we validated the structure of selected variants using fibre-FISH. Our approach has the advantage that it does not rely on a sudden change in sequence read depth for CNV detection by a HMM, which may be compromised by poor mappability of some sequence reads in the breakpoint region and assumptions about the absence of somatic variation, with the consequence that the expected copy number reflecting an integer value. However, our approach cannot detect small copy number changes because the increase or decrease in mapped reads is very small as a proportion of the total number of mapped reads at the glycophorin region.

Previous work has shown that the DUP4 variant carried by the sample HG02554 shows somatic mosaicism, leading to the suggestion that somatic mosaicism may be a feature of glycophorin structural variants (Algady et al. 2018). In this study, our fiber-FISH analyses identified no other potential somatic variants at the glycophorin locus, showing that it is not a common feature of 1000 Genomes lymphoblastoid cell lines nor of non-DUP4 variants. This suggests that somatic mosaicism is either restricted to DUP4 variants in general or restricted to the particular DUP4 sample HG02554, although a more thorough investigation of high coverage genome sequences will be needed to address this issue.

In conclusion, we identify 9 new structural variants at the glycophorin locus, characterise breakpoints and mutational mechanisms for known and novel structural variants, and show that recombination hotspot activity has influenced the nature of the structural variants observed. For some of the variants, targeted high coverage sequence using very long reads analysis will help refine some of the breakpoints. Further efforts are needed to characterise the phenotypic effects of particular variants involving gain, loss and fusion of genes

## Acknowledgements

This work was funded by SACB PhD studentships to WA and FA and Wellcome Trust grant WT098051 (F.Y. and S.L.). This research used the ALICE High Performance Computing Facility at the University of Leicester.

---

**supplementary table 1.**
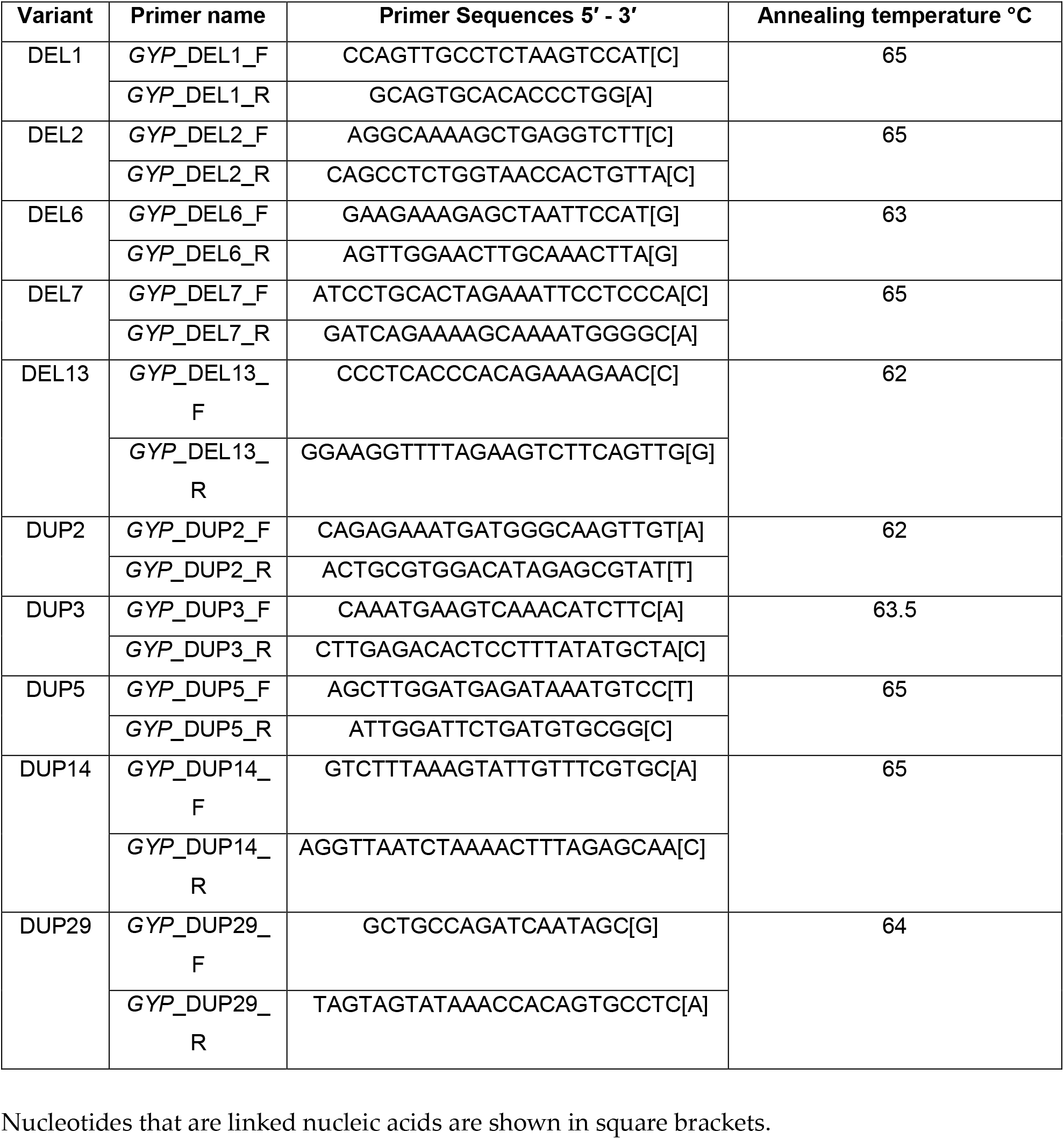

**Supplementary figure 1.**
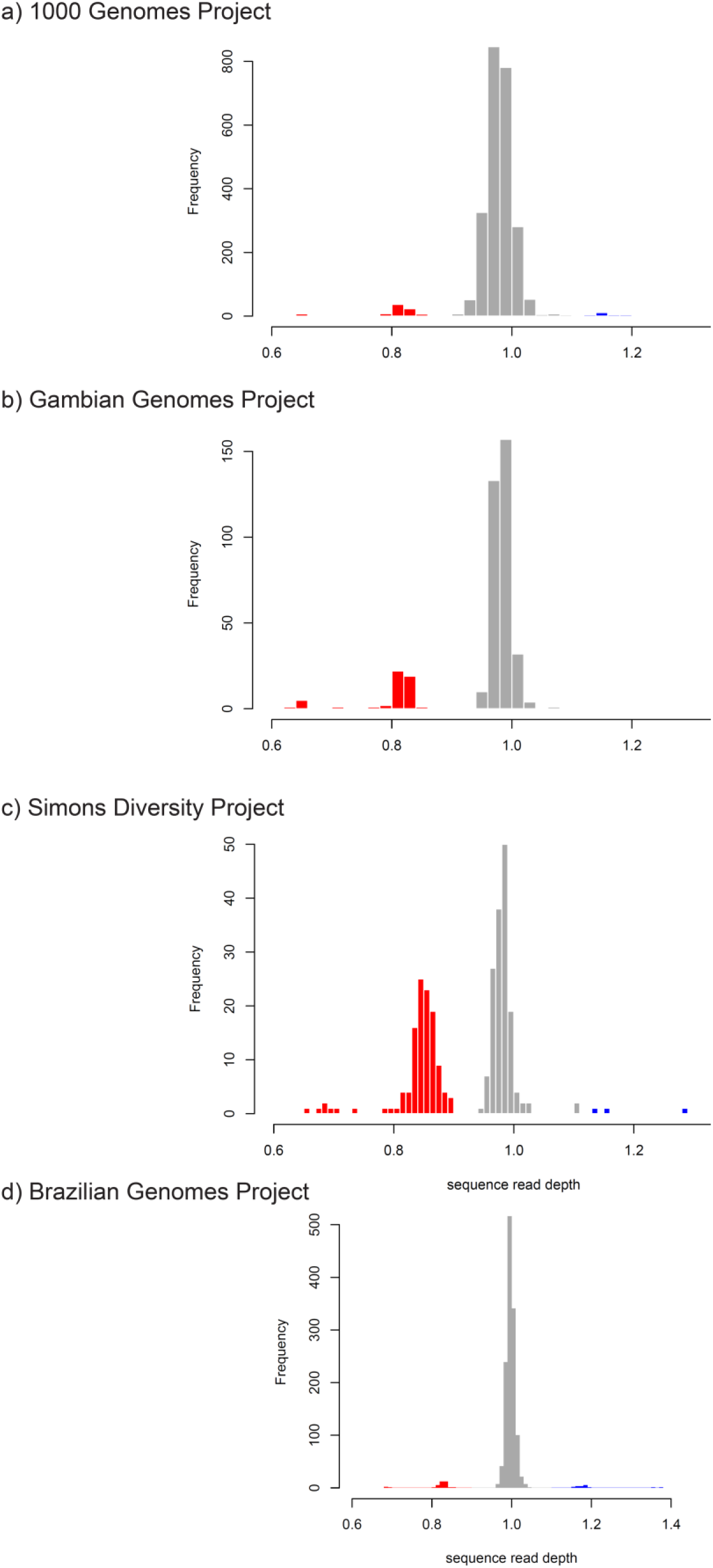
Histograms of sequence read depths of the glycophorin region. Histograms of normalised sequence read depths of the four cohorts used for this study, with red indicating putative deletions and blue putative duplications. a) 1000 Genomes Project b) Gambian Genomes Project c) Simons Diversity Project d) Brazilian Genomes Project

